# Using experience to improve: How errors shape behavior and brain activity in monkeys

**DOI:** 10.1101/203737

**Authors:** Jose L. Pardo-Vazquez, Carlos Acuña

## Abstract

Previous works have shown that neurons from the ventral premotor cortex (PMv) represent several elements of perceptual decisions. One of the most striking findings was that, after the outcome of the choice is known, neurons from PMv encode all the information necessary for evaluating the decision process. These results prompted us to suggest that this cortical area could be involved in shaping future behavior. In this work, we have characterized neuronal activity and behavioral performance as a function of the outcome of the previous trial. We found that the outcome of the immediately previous trial (*n*-1) significantly changes, in the current trial (*n*), the activity of single cells and behavioral performance. The outcome of trial *n*-2, however, does not affect either behavior or neuronal activity. Moreover, the outcome of difficult trials had a greater impact on performance and recruited more PMv neurons than the outcome of easy trials. These results give strong support to our suggestion that PMv neurons evaluate the decision process and use this information to modify future behavior.

## Introduction

The consequences of actions are fundamental for shaping future behavior. The way they are detected, represented and evaluated in the brain to guide behavior has been the focus of several research lines related to learning, in the context of value-based decisions [for reviews, see (Glimcher, 2013; Schultz & Dickinson, 2000; Wallis & Rushworth, 2013)]. In perceptual decision-making, however, the outcomes and their influence in future behavior have received less attention (Purcell & Kiani, 2016). This may be due to the fact that, once participants are trained up to their psychophysical thresholds, there is little room for learning and performance is assumed to depend mainly on sensory factors that do not change dramatically from trial to trial (Gold & Shadlen, 2007). However, it is known that, even under these circumstances, the outcomes of the preceding trials provoke behavioral adjustments, such as post-error slowing [PES; (Danielmeier & Ullsperger, 2011a; Dutilh et al., 2012; Notebaert et al., 2009; Ullsperger & Danielmeier, 2016)], post-error reduction of interference [PERI; (King, Korb, von Cramon, & Ullsperger, 2010; Ridderinkhof, 2002; Ridderinkhof et al., 2002)] and post-error improvement in accuracy [PIA; (Maier, Yeung, & Steinhauser, 2011; Marco-Pallarés, Camara, Münte, & Rodríguez-Fornells, 2008)]. The brain correlates of these adjustments have been mostly studied in humans, using EEG and imaging techniques (Danielmeier & Ullsperger, 2011b). Although these methods have provided relevant information on this topic, they lack the temporal and spatial resolution to reveal changes in the neural mechanisms of decision-making (Purcell & Kiani, 2016). These behavioral adjustments have been replicated in animal models, such as rodents (Narayanan, Cavanagh, Frank, & Laubach, 2013; Narayanan & Laubach, 2008) and non-human primates (Purcell & Kiani, 2016), also showing that their neural mechanisms seem to be well preserved across species. Therefore, the brain substrate of post-error behavioral adjustments can be studied with single cell recordings that allow a deeper understanding of the neuronal mechanisms responsible.

In perceptual tasks, single neurons from the ventral premotor cortex (PMv) represent many of the components of the decision-making process that lead to a choice, in different sensory modalities (Lemus, Hernandez, & Romo, 2009; J. L. Pardo-Vazquez, Leboran, & Acuña, 2009; Jose L Pardo-Vazquez, Leboran, & Acuna, 2008; Romo, Hermández, & Zainos, 2004). Furthermore, the firing rates of many PMv neurons are highly correlated with the behavioral choices of the animals (Acuña & Pardo-Vazquez, 2011). Remarkably, our previous work also shows that the role of these neurons does not end when the animal reaches and communicates its decision: after the behavioral response, PMv neurons represent all the information needed to evaluate the decision-process, suggesting that this area could be involved in behavioral changes resulting from such evaluation (J. L. Pardo-Vazquez et al., 2009; Jose L Pardo-Vazquez et al., 2008). If this is indeed the case, it is expected that neuronal responses during a given trial will carry information of the preceding outcomes. In this work, we have tested this prediction by recording single cells from PMv while one monkey performed a visual discrimination task and characterizing the effects of previous outcomes on neuronal activity and behavioral performance.

We found that 27% of the recorded neurons show different firing rates, during trial the current trial (*n*), after correct and error choices in immediately preceding trial (*n*-1). Behavioral performance was also different in post-correct and post-error trials: the monkey made less aborts after errors (post-error behavioral engagement) and improved its accuracy (post-error behavioral accuracy). There was no effect whatsoever when neural activity and behavior were conditioned on the outcomes of trial *n*-2. Finally, the effect of the previous outcome was stronger, both at the behavioral and neuronal level, when the previous trial was difficult than when it was easy.

## Materials and Methods

**General.** Experiments were carried out on one male monkey (Macaca mulatta). The animal (BM7, 8 kg) was handled according to the standards of the European Union (86/609/EU), Spain (RD 1201/2005) and the SFN Policies and Use of Animals and Humans in Neuroscience Research. The experimental procedures were approved by the Bioethics Commission of the University of Santiago de Compostela (Spain). The monkey’s head was fixed during the task and looked binocularly at a monitor screen placed at 114 cm away from their eyes (1 cm subtended 0.5° to the eye). The room was isolated and soundproof. Two circles (1° in diameter) were horizontally displayed 6° at the right and 6° at the left of the fixation bar (a vertical bar; 0.5° length, 0.02° wide) displayed in the center of the screen. The visual stimuli –lines with different lengths-were displayed in the center of the screen. The monkey used right and left circles to indicate, with an eye movement, whether the second stimulus (S2) was shorter or longer than the first stimulus (S1), respectively. Binocular eye movements were recorded with SMI iView X Hi-Speed Primate, sampled at 500 Hz and acquired with MonkeyLogic 1.0 (www.monkeylogic.org; (Asaad & Eskandar, 2008). The eye position was calibrated daily using the five points automatic routine included in this toolbox. Visual stimuli were created in a 2.67 GHz Intel^®^ Core^™^ i7 PC using a 1024 MB NVIDIA GeForce GT 240 graphic card and presented in an ASUS VH226H monitor, with 60 Hz vertical refresh rate. The monitor mode when the task was running was 1920 × 1080 (75 Hz). MonkeyLogic 1.0 was used for task control and to generate visual stimuli. The experiment consisted of two phases. The first one (training phase) lasted for about 12 months and was aimed at training the monkey and estimating the psychometric functions relating performance (proportion choices “longer than”) with the difference in length between S1 and S2. These functions were then used to select a reduced set of stimuli for the second phase (recording phase), based on the discrimination capability of the monkey. Single cell recordings were performed during the second phase.

**Stimuli.** The visual stimuli were stationary bright lines, subtending 0.15° in width. During the training phase, three different lengths (2, 2.18 and 2.36°) were used as S1. For each of them, 10 lengths (five longer and five shorter) were used as S2, in 0.09° steps. This stimuli set allowed us to confirm that the animal was using the difference in length between S2 and S1 to solve the task and to reliably estimate the psychometric functions relating this difference with the probability of perceiving S2 as “longer than” S1. These functions were then used to select the stimuli set for the recording phase depending on the monkey’s capability to discriminate. In this second phase, we used the same three lengths as S1 and four lengths (two longer and two shorter) as S2 for each S1. The lengths of the S2 were selected, independently for S2 shorter than S1 and S2 longer than S1, so as to provoke 90 and 70% correct discriminations, for easy and difficult conditions, respectively.

**Discrimination Task.** The monkey was trained for about one year to discriminate up to its psychophysical threshold in a 2-alternative forced-choice task, the length discrimination task (LDT) sketched and explained in Figure 1A. The lengths of S1 and S2 changed randomly from trial to trial and aborts were repeated. The inter-trial interval was 1500 ms. During the training phase, in which 30 conditions were presented, trials were grouped in blocks of 240 trials. During the recording phase, in which only 12 conditions were presented, trials were grouped in blocks of 192 trials.

**Figure 1.**
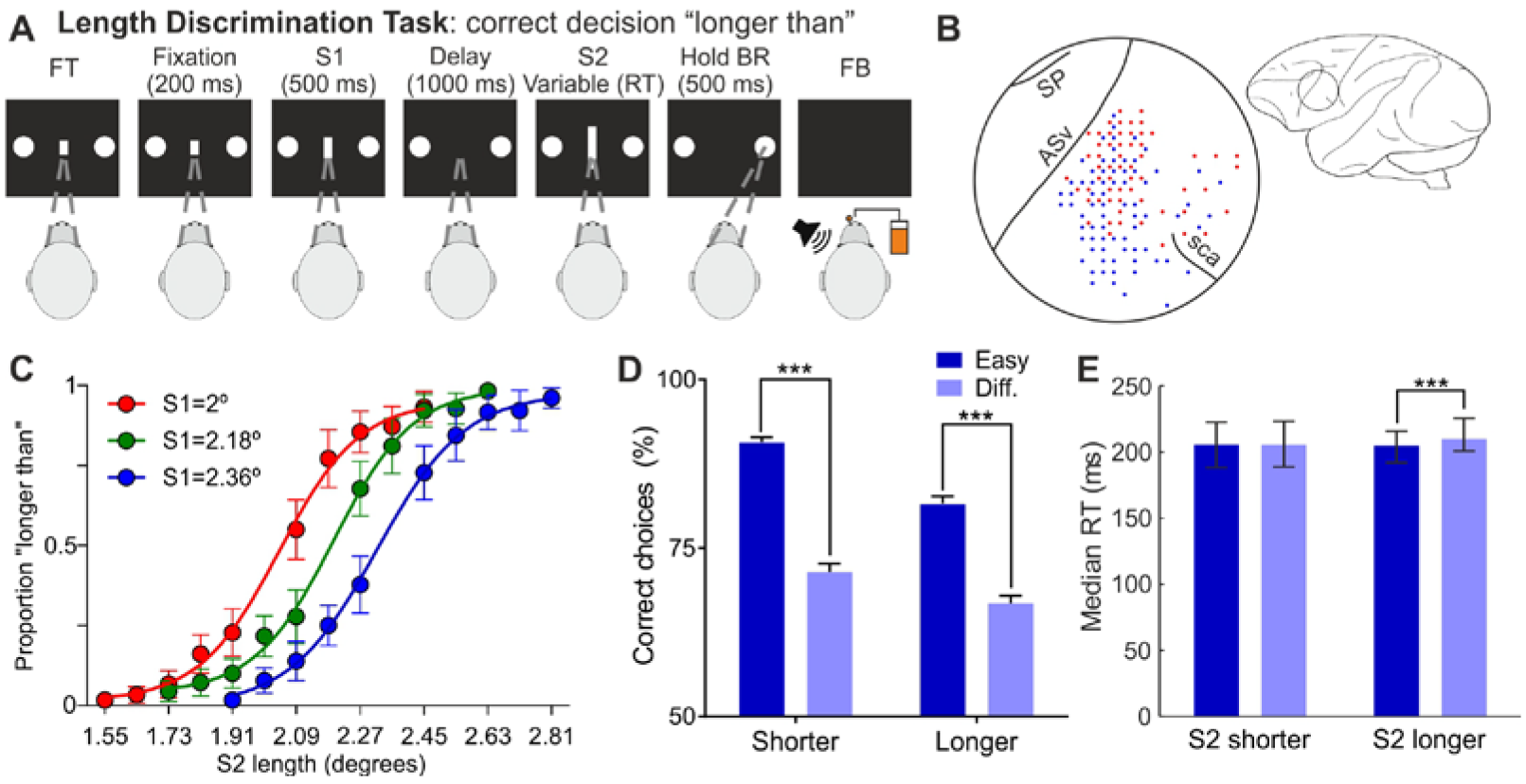
Behavioral task, behavioral performance and recordings localization. **(A)** Sequence of events during the length discrimination task. The fixation target (FT) and two response buttons appear on screen. The monkey initiates the trial by fixating in the FT and, after a 200 ms fixation time, two lines of variable length (S1 and S2) are presented sequentially, separated by a 1 second delay. S1 is presented for 500 ms and S2 remains on screen until the subject indicates, with an eye movement, whether S2 is longer (right button) or shorter (left button) than S1. After the behavioral response (BR), the monkey has to hold the response for 500 ms and then feedback (FB) is provided. Correct choices are indicated with a 60 dB SPL, high pitch (3000 Hz) sound (200 ms) and a drop of liquid; incorrect choices are indicated with a 60 dB SPL, low pitch (500 Hz) sound (200 ms). The inter-trial interval is 1,500 ms. **(B)** Localization of the recordings. A total of 658 neurons, recorded in 146 penetrations, were analyzed. Red and blue dots indicate the localization of the penetrations performed in the left (n=60) and right (n=86) hemispheres, respectively. SP, *sulcus principalis*; ASv, *arcuate sulcus ventralis*; sca, *sulcus subcentralis anterior*. **(C)** Psychometric functions, one per S1 length, estimated in the end of the training phase. Dots represent proportions averaged across 30 blocks of trials; error bars represent 95% CI. **(D, E)** Behavioral accuracy (mean ±95% CI) and RTs (median ± 1 quartile), respectively, during the recording sessions, showing the effect of difficulty on discrimination accuracy and reaction times. ***, p<0.001.

**Recordings.** Extracellular single unit activity was recorded with tungsten microelectrodes (1.5-3.5 MΩ) in the posterior bank of the ventral arm of the sulcus arcuatus and adjacent surface in the ventral premotor cortex in the two hemispheres of the monkey. Standard histological techniques were used to confirm the location of the penetrations (Figure 1B). Recording sites changed from session to session.

**Data analysis**. All analyses were carried out using custom-made programs in Matlab 2012b (http://www.mathworks.com). Firing rates were estimated by counting the number of spikes within 10 ms bins and then averaging them with a 200 ms sliding window (10 ms steps). The effect of previous outcomes on neuronal activity was assessed with Receiver Operating Characteristics [ROC; (Green & Swets, 1966)] analysis, which allows the measure of the degree of overlap between two response distributions. For each neuron with at least 10 correct and 10 incorrect trials, we computed the area under the ROC curve (AUC ROC) within a 200 ms bin that was slid in 10 ms steps [see (J. L. Pardo-Vazquez et al., 2009; Jose L Pardo-Vazquez et al., 2008) for detailed information on this method]. AUC ROC values significance was assessed with a permutation test (*n*=2000 iterations), significance level was set at *p*<0.01, corrected for multiple comparisons (Fujisawa, Amarasingham, Harrison, & Buzsáki, 2008). Psychometric curves were estimated with the same procedure described elsewhere (J. L. Pardo-Vazquez et al., 2009; Jose L Pardo-Vazquez et al., 2008). Significance of behavioral comparisons was evaluated with a permutation test (n=20000 iterations).

## Results

The activity of single cells was recorded while one monkey performed a length discrimination task (LDT; Figure 1A). A total of 146 penetrations (60 and 86 in the left and right hemispheres respectively) were performed in the PMv (Figure 1B). In the task, two lines (S1 and S2) were presented sequentially and separated by a 1 s delay, and the monkey had to decide whether S2 was longer or shorter than S1 and communicate its decisions with an eye movement.

### Psychometric functions, stimulus selection and behavioral performance during the recordings

During the training phase, three S1 and ten S2 per S1 (see Methods) were used to estimate the psychometric functions relating the proportion of responses “longer than” with the length of S2 (Figure 1C). These functions, which suggest that the subject based its choices on the difference between S2 and S1, were then used to select the stimuli set for the recording phase: S2 lengths that provoked 90 and 70 correct choices, for easy and difficult conditions respectively, were independently chosen for each S1 and for “shorter than” (S2<S1) and “longer than” (S2>S1) conditions. For the 2° S1, the S2 could be 1.78° and 2.35° for easy conditions and 1.95° and 2.16 for difficult conditions. For the 2.18° S1, the S2 could be 1.9° and 2.45° for easy conditions and 2.08° and 2.29° for difficult conditions. Finally, for the 2.36° S1, the S2 could be 2.06° and 2.61° for easy conditions and 2.23° and 2.44° for difficult conditions.

Performance during the recording sessions (Figure 1D) shows that accuracy in the LDT depends on the difficulty of the discrimination and that the monkey was using the lengths of S2 and S1 to solve the task also for this subset of conditions. For S2<S1 trials, the percentages of correct responses were 91 and 71% for easy and difficult conditions, respectively. For S2>S1 trials, the percentage of correct responses were 82 and 67%. The differences between easy and difficult trials were significant for both S2<S1 and S2>S1 discriminations (p<0.001). Note that the average percentage of correct responses closely matches the target (90 and 70%) we established for selecting the stimuli set.

Regarding discrimination speed (Figure 1E), task difficulty had little effect on reaction times (RTs), that were slightly and significantly faster only for easy S2>S1 discriminations. For correct S2>S1 trials, the median RTs were 210 ms (1st Q = 201 ms; 3rd Q = 226 ms) and 205 ms (1st Q = 192; 3rd Q = 216 ms) for difficult and easy discriminations, respectively, and they were significantly different (*p*<0.001). For correct S2<S1 trials, the median RTs were 205 ms (1st Q = 188; 3rd Q = 223 ms) and 206 ms (1st Q = 189 ms; 3rd Q = 223 ms) for difficult and easy discriminations, respectively, and they were not significantly different (*p*=0.91).

### *The outcome of the previous trial (*n-*1) has significant effects on behavior and neuronal activity during the current trial* (n)

Since we were interested in proving the contribution of PMv neurons to shaping behavior based on the outcomes of previous trials, the first step was to verify whether these outcomes had significant effects on performance in the LDT. To this end, we analyzed all the data gathered during the recording sessions (88,400 trials divided in 442 blocks of 200 trials). Behavioral accuracy was calculated as the mean percentage of correct choices with respect to the number of completed trials, for the subsets of trials preceded by errors and correct choices separately. We found a significant (*p*<0.01), post-error improvement in accuracy (PIA), from 76.9% (*SD* = 5.5%) correct decisions after correct trials to 78.5% (*SD* = 9%) after errors (Figure 2A).

**Figure 2.**
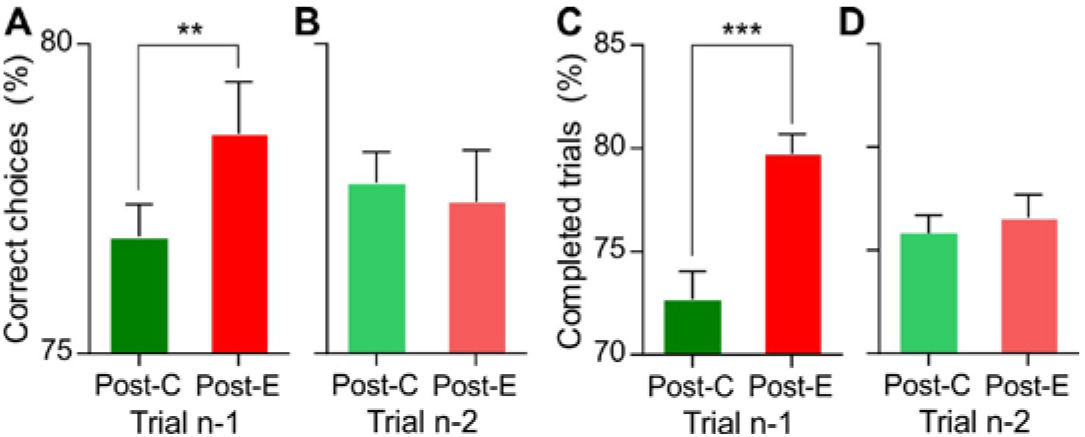
Behavioral performance on trial *n* is affected by the outcome of trial *n*-1, while the outcome of trial *n*-2 has no significant effect. **(A, B)** Behavioral accuracy as a function of the outcome of the previous trials; proportion of correct and incorrect trials over the number of completed trials conditioned on the outcome of trials *n*-1 and *n*-2, respectively. **(C, D)** Behavioral engagement as a function of the outcome of the previous trials; proportion of completed trials conditioned on the outcome of trials *n*-1 and *n*-2, respectively. Bars represent mean ± 95% CI; ***, *p*<0.001; **, *p*<0.01.

Since, in perceptual tasks, performance is mainly limited by sensory constrains that are not expected to change dramatically from trial to trial, we looked for changes in behavioral engagement, as reflected in an alteration of the percentages of completed trials. We found that the outcome of the preceding choice produced a significant posterror improvement in engagement (PIE; Figure 2C): the percentage of completed trials was significantly higher (*p*]<0.001) in post-error (79.7%, *SD*=10.1%) than in post-correct trials (72.7%, *SD*=14.3%).

The outcome of preceding trials is also known to affect speed; to compare the RTs following correct choices and errors, we split the 88,400 trials in 442 blocks of 200 trials, estimated the median RT after correct and error trials for each block and then compared their means. We found the RTs to be slightly, but significantly, faster (*p*<0.001) in post-error trials (mean = 205 ms, *SD* = 8.9 ms) than in post-correct trials (mean = 207 ms, *SD* = 7.2 ms).

Therefore, our behavioral results confirm that, in the LDT, the outcome of the previous trial had a significant effect on performance in the current trial, increasing both the accuracy of the discriminations and also the probability for the monkey to be engaged in the task and complete the trial.

To assess whether neuronal activity during the current trial (*n*) reflects the outcome of the previous trial (*n*-1), we compared the firing rate of single cells in post-correct (Post-C) and post-error (Post-E) trials and used ROC analyses to quantify that effect. We focused the analysis in the time period going from the onset of S1 to the end of the delay, to avoid differential neuronal responses owed to the decision process in the current trial. Figure 3 shows the activity of two example neurons during trial *n*, conditioned on the outcome of trial *n*-1. One of them shows higher firing rates after incorrect trials (Post-C<Post-E; Figure 3A and C) and the other responds more strongly after correct trials (Post-C>Post-E; Figure 3B and D). ROC analysis confirmed, for both neurons, the existence of a trial-by-trial representation of the outcome of the preceding choice in the activity of the current trial (Figure 3E and F).

**Figure 3.**
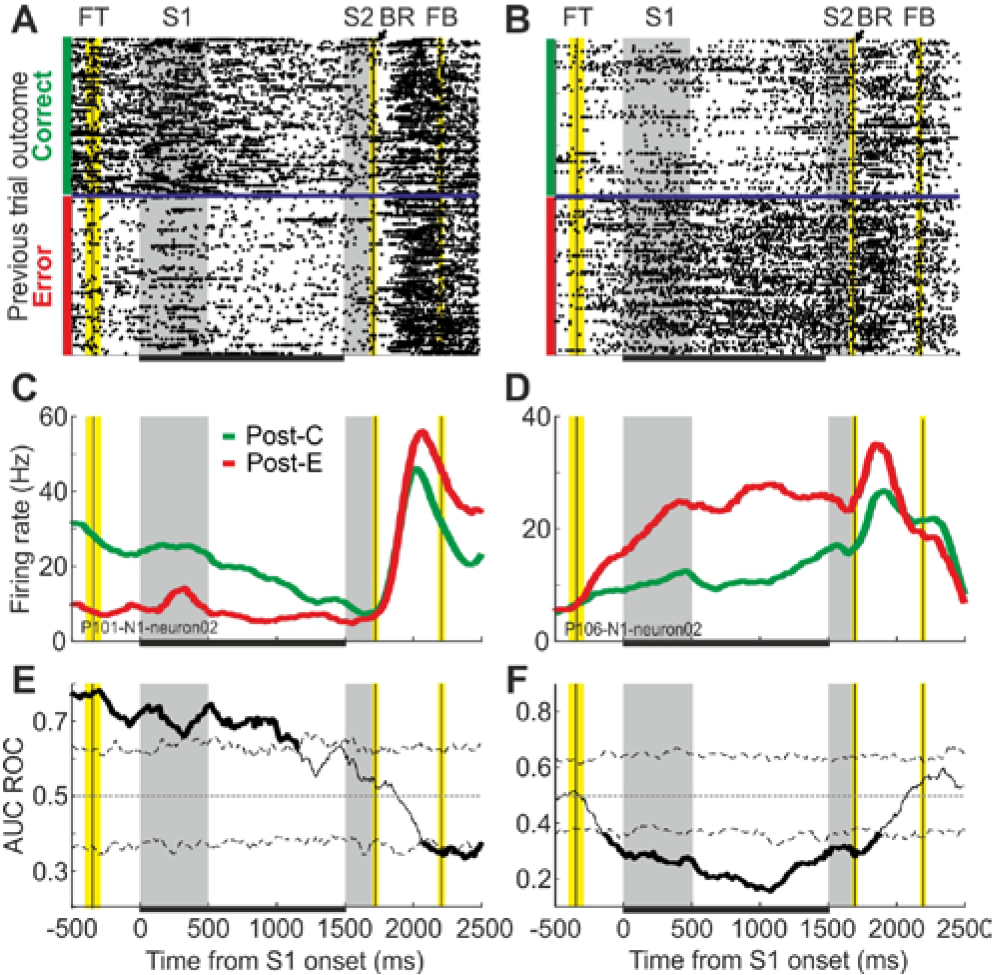
Single cells show significantly different responses in a given trial (*n*) depending on the outcome of the previous trial (*n*-1). **(A, B)** Raster plots, sorted as a function of the outcome of the previous trial (*n*-1), for two example neurons showing higher firing rates after correct and after incorrect choices, respectively. Horizontal blue lines separate trials preceded by correct and incorrect choices. Grey shaded areas indicate S1 and S2 presentation time; vertical black lines and yellow shaded areas under the FT (fixation target), BR (behavioral response) and FB (feedback) labels indicate the average time of these task events ± SD; horizontal black bars in the bottom indicate the relevant period analyzed here. **(C, D)** Firing rate averaged across post-correct (Post-C) and post-error (Post-E) trials, for the same neurons in A and B. Firing rates were estimated using a sliding window (200 ms length, 10 ms steps). **(E, F)** Area under the ROC curve (AUC ROC) comparing post-correct and post-error trials. Significance thresholds (dashed lines) were estimated with a permutation test (n=2000). The analysis was conducted with a sliding window (200 ms, 10 ms steps); α=0.01; corrected for multiple comparisons.

Out of 658 neurons with enough trials to be analyzed (see Methods), 175 (27%) showed significant AUC ROC values (*p*<0.01) within the relevant period, *i.e*., at least one significant bin during the first 1500 ms of the trial (Figure 4A). Out of these 175 neurons, 104 (59%) showed higher firing rates after errors and 71 after correct choices. Figure 4C shows the AUC ROC averaged across Post-C<Post-E and Post-C>Post-E populations of neurons; the joint activity of these populations represents, during the whole duration of the current trial, the outcome of the previous choice. Figure 4E and F show the localization of these neurons in the cortex and the distribution of minima and maxima significant AUC ROC values, respectively; the mean for Post-C<Post-E and Post-C>Post-E neurons were 0.31 (*SD* = 0.07) and 0.67 (*SD* = 0.06), respectively.

**Figure 4.**
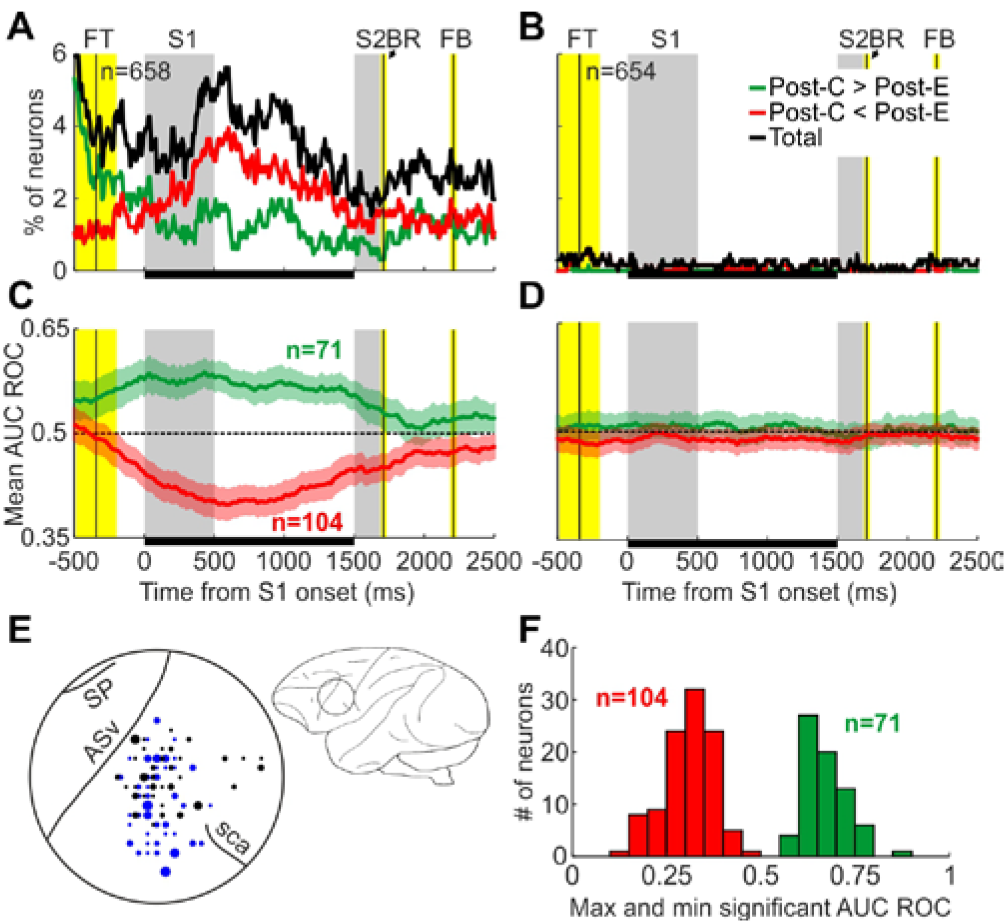
Neuronal activity shows the same pattern as behavioral performance: the outcome of the previous trial (*n*-1) has significant effects on neuronal activity while the outcome of trial *n*-2 does not affect it. **(A, B)** Percentage of neurons showing significantly different responses on trial n as a function of the outcome (correct or error) of trials *n*-1 and *n*-2, respectively. Green and red lines represent neurons preferring post-correct (Post-C>Post-E) and post-error (Post-C<Post-E) trials, respectively. **(C)** Area under the ROC curve averaged across the populations of Post-C>Post-E (green) and Post-C<Post-E (red) neurons. These populations were established by selecting those neurons with significant AUC ROC values between 0 and 1500 ms from stimulus onset. Shaded areas indicate mean ± 2.5 SEM. **(D)** Area under the ROC curve averaged across the same populations in C, but conditioned on the outcome of trial *n*-2. **(E)** Localization of neurons in the PMv with significant AUC ROC values between 0 and 1500 ms from S1 onset. Black and blue dots represent neurons recorded in the left and right hemispheres, respectively; dot size indicates the number of neurons, from 1 to 7. **(F)** Distribution of maxima and minima significant AUC ROC values for Post-C>Post-E (green) and Post-C<Post-E (red) neurons.

### *The outcome of trial* n-*2 does not affect either behavior or neuronal activity in the current trial* n)

We assessed the duration of post-error improvements by repeating the same analyses but conditioned on the outcome of trial *n*-2. When the percentage of correct choices over the total number of completed trials-behavioral accuracy-was compared (Figure 2B), no significant difference (*p*=0.53) was observed as a function of the outcome of the previous trial (*n*-2): the mean percentages of correct choices were 77.8% (*SD*=5.3%) and 77.4% (*SD*=8.9%) after correct choices and errors, respectively.

Behavioral engagement was also very similar for post-correct (*n*-2) and post-error (*n*-2) trials (Figure 2D): the percentage of completed trials after correct choices in the trial *n*-2 (75.8%; *SD* = 9.3%) was not different (*p*=0.298) from the percentage after errors in the trial *n*-2 (76.6%; *SD*=12%).

Remarkably, we found the same pattern when analyzing the effect of the outcome of trial *n*-2 on neuronal activity: less than 5% of the neurons (26 out of 654 neurons analyzed) reached significant AUC ROC values (Figure 4B) and the average AUC-ROCs for post-correct and post-error preferring neurons were very close to 0.5 (Figure 4D). Therefore, while the outcome of trial *n*-1 had a significant impact on the activity of PMv neurons and behavioral accuracy and engagement, the outcome of trial *n*-2 had a much smaller effect on the neuronal activity and no significant effect on behavioral performance.

### *During the current trial (*n*), the outcome of difficult trials (*n-*1) has stronger effects on behavior and recruits more PMv neurons than the outcome of easy trials*

To study the combined effect of the outcome and the difficulty of the previous trial, we analyzed outcome-related changes in behavior and neuronal activity after easy and difficult choices separately. The difficulty of the previous trial (*n*-1) had significant effects on behavioral adjustments.

On the one hand, we only found a significant PIA after difficult trials (Figure 5A). For post-easy trials, the mean percentages of correct responses after correct and error trials were 76.9% (*SD* = 7.6%) and 78.3% (*SD* = 16.7%), respectively, and this difference was not significant (*p*=0.11). For post-difficult trials, the mean percentages of correct responses after correct and error trials were 76.7%(*SD* = 9.1%) and 79% (*SD* = 11%), and this difference was significant (*p*<0.01).

**Figure 5.**
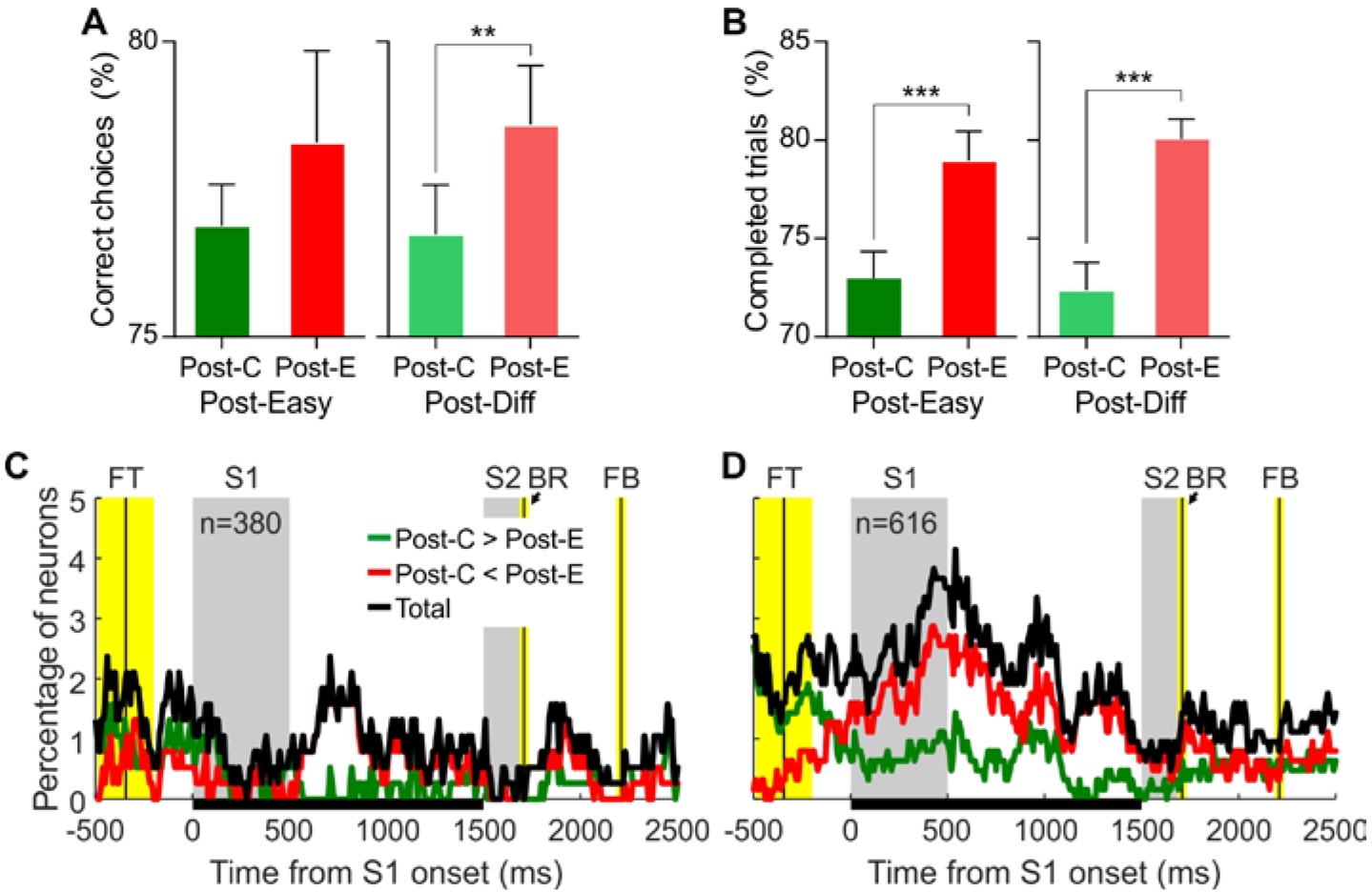
The outcome of difficult trials has a stronger effect on behavioral performance and neural activity than that of easy trials. **(A)** Combined effect of outcome and difficulty of the previous trial (*n*-1) on behavioral accuracy. **(B)** Combined effect of outcome and difficulty of the previous trial (*n*-1) on behavioral engagement. Bars represent mean ± 95% CI; ***, *p*<0.001; **, *p*<0.01. **(C, D)** Percentage of neurons showing significant effect of the outcome of the previous choice (*n*-1) following easy and difficult trials, respectively.

On the other hand, although PIE was significant both after easy and difficult trials, it was higher in the latter case (Figure 5B). For post-easy trials, the mean percentages of completed trials after correct and error trials were 73% (*SD*=14.7%) and 79% (*SD*=16.1%), respectively. For post-difficult trials, these percentages were 72.4% (*SD*=15%) and 80.1% (*SD*=10.9%), respectively. Both differences were significant (*p*<0.001). Together, these results show that behavioral improvement after errors was higher when those errors happened in difficult trials.

Regarding neuronal activity, when only easy trials were considered, 48 (13%) neurons out of the 380 with enough trials showed significant AUC ROC values (Figure 5C). From these, 28 (58%) were Post-C<Post-E neurons and 20 were Post-C>Post-E. When only difficult trials were considered, 103 (17%) neurons out of the 616 with enough trials showed significant AUC ROC values (Figure 5D). From these, 66 (64%) were Post-C<Post-E neurons and 37 were Post-C>Post-E. Furthermore, the duration of the effect of the outcome of the previous trial on the current was also different as a function of the difficulty of the trial *n*-1: the average number of significant bins after easy and difficult discriminations was 11 (*SD* = 16.4) and 20 (*SD* =28.6), respectively. The temporal profile of the neuron count also shows this difference: during the 1500 ms in which our analysis was focused, the percentage of recruited neurons is clearly higher throughout the trial duration (Figure 5C and D). Therefore, although there are PMv neurons representing the outcome of the previous choice (*n*-1) both after easy and difficult trials, our results suggest that this effect is higher in the latter case, as it happened with behavior.

## Discussion

In this work, we show, for the first time, that single neurons from PMv represent, during the current trial, the outcome of the immediately previous one. Post-correct and post-incorrect firing rates are significantly different in 27% of the analyzed neurons. Behavioral results demonstrate that the outcome of the preceding trial affects performance in a visual discrimination task, increasing accuracy and engagement in the task. These changes in neuronal activity and behavior during the current trial are only provoked by the outcome of the immediately previous trial. Finally, when the difficulty of the preceding trial was considered, we found that the effect of the outcome is stronger after difficult trials, again at both the neuronal and behavioral levels.

Post-error behavior in the LDT is consistent with previous works and also shows a new effect, post-error increase in engagement, which should be considered when using animal models for addressing behavioral adjustments resulting from the consequences of previous actions. Post-error slowing has been shown, mostly in humans, in a variety of behavioral tasks (Danielmeier & Ullsperger, 2011a; Debener et al., 2005; Dutilh et al., 2012; King et al., 2010; Notebaert et al., 2009); it has also been described in monkeys performing the random dots motion (RDM) task (Purcell & Kiani, 2016) and in rats performing a time estimation task (Narayanan & Laubach, 2008). The phenomenology of PES is complex and it seems to depend on many task variables (Danielmeier & Ullsperger, 2011a; Dutilh et al., 2012; Notebaert et al., 2009), such as inter-trial interval and outcome frequency. In fact, when correct trials are infrequent, RTs are faster after error trials (Castellar, Kühn, Fias, & Notebaert, 2010; Notebaert et al., 2009). In the current work, we found a weak but significant post-correct speeding that can be explained by two features of the LDT: first, unlike in the RDM task, S1 and S2 do not change in time and the monkey does not get a clear benefit from increasing the sampling time –as suggested by the effect of the difficulty on the RTs–; and second, performance in the LDT does not depend only in correctly perceiving the length of S2, but also on perceiving and maintaining in working memory the length of S1. Therefore, increasing the RT after errors would not have a great impact on accuracy. Despite the abundance of research focused on understanding the functional meaning of PES, it is still under debate whether this behavioral adjustment has an adaptive role, *i.e*., whether it translates into better chances of correctly solving post-error trials (Danielmeier & Ullsperger, 2011a; Notebaert et al., 2009; Ullsperger & Danielmeier, 2016; Van der Borght, Desmet, & Notebaert, 2016; Van Der Borght, Schevernels, Burle, & Notebaert, 2016). Together with others (Purcell & Kiani, 2016), our results might help shedding light on this debate. On the one hand, in tasks in which there is an advantage in sampling the stimulus for a longer time, animals slow down their decisions after errors (Purcell & Kiani, 2016). On the other hand, when the behavioral task is designed so that sampling for longer time does not benefit the decision, as in the LDT, the subject does not show slower RTs after errors. Therefore, it seems that animals only slow down the decision, after an error, when this strategy can be beneficial to them.

Regarding post-error improvement in accuracy, it has been previously described in humans, mostly using attentional paradigms such as the flanker task (Maier et al., 2011; Marco-Pallarés et al., 2008). Other experiments, however, did not find differences in accuracy as a function of the outcome of the preceding trial (Hajcak, McDonald, & Simons, 2003; Hajcak & Simons, 2008; King et al., 2010) and in some cases even decreased accuracy after errors was reported (Castellar et al., 2010; Notebaert et al., 2009). In monkeys (Purcell & Kiani, 2016) and rodents (Narayanan & Laubach, 2008), errors did affect RTs, but no PIA was reported. Although significant, the size of PIA in the current work was modest (less than 2%), as expected given the nature of our task. Perceptual decisions depend mostly on sensory processing and dramatic changes in discrimination thresholds are not to be expected from one trial to the next. Therefore, in subjects trained up to their psychophysical thresholds, there is little room for improving performance after an error is detected.

This is the first time, to our knowledge, that post-error improvement in engagement is described. This effect could result from an increase in motivation and attentional resources devoted to the task to avoid further errors (Maier et al., 2011). It could also explain faster post-error RTs in the LDT, as a consequence of the monkey being highly motivated and engaged in the task after making an error. Such a behavioral adjustment is hard to find in humans, since they typically do not fail to complete many trials. However, PIE might be relevant when studying the effects of the outcomes of previous trials and their neural basis in rodents and non-human primates, which usually abort a significant percentage of the trials (*e.g*., more than 20% of the trials in the LDT).

Differential neuronal responses after correct and error trials have been described in the dorsomedial prefrontal cortex (dmPFC) of rats performing a time estimation task (Narayanan & Laubach, 2008), providing further evidence of the involvement of this area in post-error behavioral adjustments, since its inactivation attenuated PES. In dmPFC, two populations of neurons were described, one with sustained increased firing rates after incorrect trials and the other after correct trials. These results were interpreted as evidence of a form of retrospective memory aimed at monitoring task performance. The neuronal responses described in dmPFC of rats are very similar to the ones we found, in the present work, in monkey’s PMv.

Remarkably, the effect of the consequences of the previous trials on neuronal response in the PMv mimicked post-error behavioral adjustments. Firstly, while the outcome of trial *n*-1 showed significant effects on discrimination performance and neuronal activity in trial *n*, the outcome of trial *n*-2 showed no effect on either of them. Secondly, behavioral and neuronal effects were stronger after difficult trials as compared to easy ones. These results provide support to our claim that PMv could be involved in using the consequences of previous actions to shape future behavior, most likely as part of a broader network of decision-related brain regions, including parietal (Purcell & Kiani, 2016) and prefrontal (Narayanan & Laubach, 2008) cortices. Further research, using methods such as electrical stimulation or reversible inactivation to manipulate neural activity in this area, should be conducted in order to stablish a causal relationship between neuronal activity in PMv and post-error behavioral adjustments.

## Acknowledgements

This research has been supported by the following grants: Human Frontier Science Program Long-Term Award (LT00042/2012) to Jose L. Pardo-Vazquez; from Ministerio de Ciencia e Innovatión (MICINN), Spain, to Carlos Acuña.

## References

Acuna, C., & Pardo-Vazquez, J. L. (2011). Ventral premotor cortex neuronal activity matches perceptual decisions. European Journal of Neuroscience, 33(12), 2338–2348. http://doi.org/10.1111/j.1460-9568.2011.07708.x

Asaad, W. F., & Eskandar, E. N. (2008). Achieving behavioral control with millisecond resolution in a high-level programming environment. Journal of Neuroscience Methods, 173(2), 235–240. http://doi.org/10.1016/j.jneumeth.2008.06.003

Castellar, E. núňez, Kühn, S., Fias, W., & Notebaert, W. (2010). Outcome expectancy and not accuracy determines posterror slowing: ERP support. Cognitive, Affective, & Behavioral Neuroscience, 10(2), 270–278. http://doi.org/10.3758/CABN.10.2.270

Danielmeier, C., & Ullsperger, M. (2011a). Post-Error Adjustments. Frontiers in Psychology, 2, 233. http://doi.org/10.3389/fpsyg.2011.00233

Danielmeier, C., & Ullsperger, M. (2011b). Post-error adjustments. Frontiers in Psychology, 2, 233. http://doi.org/10.3389/fpsyg.2011.00233

Debener, S., Ullsperger, M., Siegel, M., Fiehler, K., Cramon, D. Y. von, & Engel, A. K. (2005). Trial-by-Trial Coupling of Concurrent Electroencephalogram and Functional Magnetic Resonance Imaging Identifies the Dynamics of Performance Monitoring. The Journal of Neuroscience, 25(50), 11730–11737. http://doi.org/10.1523/JNEUROSCI.3286-05.2005

Dutilh, G., Vandekerckhove, J., Forstmann, B. U., Keuleers, E., Brysbaert, M., & Wagenmakers, E.-J. (2012). Testing theories of post-error slowing. Attention, Perception, & Psychophysics, 74(2), 454–465. http://doi.org/10.3758/s13414-011-0243-2

Fujisawa, S., Amarasingham, A., Harrison, M. T., & Buzsáki, G. (2008). Behavior-dependent short-term assembly dynamics in the medial prefrontal cortex. Nature Neuroscience, 11(7), 823–833. http://doi.org/10.1038/nn.2134

Glimcher, P. W. (2013). Value-Based Decision Making. In Neuroeconomics: Decision Making and the Brain: Second Edition (pp. 373–391). http://doi.org/10.1016/B978-0-12-416008-8.00020-6

Gold, J. I., & Shadlen, M. N. (2007). The neural basis of decision making. Annual Review of Neuroscience, 30, 535–74. http://doi.org/10.1146/annurev.neuro.29.051605.113038

Green, D. M., & Swets, J. A. (1966). Signal detection theory and psychophysics. New York: Wiley.

Hajcak, G., McDonald, N., & Simons, R. F. (2003). To err is autonomic: Error-related brain potentials, ANS activity, and post-error compensatory behavior. In Psychophysiology (Vol. 40, pp. 895–903). http://doi.org/10.1111/1469-8986.00107

Hajcak, G., & Simons, R. F. (2008). Oops!.. I did it again: An ERP and behavioral study of double-errors. Brain and Cognition, 68(1), 15–21. http://doi.org/10.1016/j.bandc.2008.02.118

King, J. A., Korb, F. M., von Cramon, D. Y., & Ullsperger, M. (2010). Post-Error Behavioral Adjustments Are Facilitated by Activation and Suppression of Task-Relevant and Task-Irrelevant Information Processing. Journal of Neuroscience, 30(38), 12759–12769. http://doi.org/10.1523/JNEUROSCI.3274-10.2010

Lemus, L., Hernandez, a, & Romo, R. (2009). Neural encoding of auditory discrimination in ventral premotor cortex. Proc Natl Acad Sci U S A, 106(34), 14640–14645. http://doi.org/10.1073/pnas.0907505106

Maier, M. E., Yeung, N., & Steinhauser, M. (2011). Error-related brain activity and adjustments of selective attention following errors. NeuroImage, 56(4), 2339–2347. http://doi.org/10.1016/j.neuroimage.2011.03.083

Marco-Pallarés, J., Camara, E., Münte, T. F., & Rodríguez-Fornells, A. (2008). Neural mechanisms underlying adaptive actions after slips. Journal of Cognitive Neuroscience, 20(9), 1595–610. http://doi.org/10.1162/jocn.2008.20117

Narayanan, N. S., Cavanagh, J. F., Frank, M. J., & Laubach, M. (2013). Common medial frontal mechanisms of adaptive control in humans and rodents. Nature Publishing Group, 16(12), 1888–1895. http://doi.org/10.1038/nn.3549

Narayanan, N. S., & Laubach, M. (2008). Neuronal Correlates of Post-Error Slowing in the Rat Dorsomedial Prefrontal Cortex. Journal of Neurophysiology, 100(1), 520–525. http://doi.org/10.1152/jn.00035.2008

Notebaert, W., Houtman, F., Opstal, F. Van, Gevers, W., Fias, W., & Verguts, T. (2009). Post-error slowing: An orienting account. Cognition, 111(2), 275–279. http://doi.org/10.1016/j.cognition.2009.02.002

Pardo-Vazquez, J. L., Leboran, V., & Acuna, C. (2009). A role for the ventral premotor cortex beyond performance monitoring. Proceedings of the National Academy of Sciences, 106(44), 18815–18819. http://doi.org/10.1073/pnas.0910524106

Pardo-Vazquez, J. L., Leboran, V., & Acuña, C. (2008). Neural correlates of decisions and their outcomes in the ventral premotor cortex. Journal of Neuroscience, 28(47), 12396–12408. Retrieved from http://www.ncbi.nlm.nih.gov/pubmed/19020032

Purcell, B. A., & Kiani, R. (2016). Neural Mechanisms of Post-error Adjustments of Decision Policy in Parietal Cortex. Neuron, 89(3), 658–671. http://doi.org/10.1016/j.neuron.2015.12.027

Ridderinkhof, K. R. (2002). Micro-and macro-adjustments of task set: Activation and suppression in conflict tasks. Psychological Research, 66(4), 312–323. http://doi.org/10.1007/s00426-002-0104-7

Ridderinkhof, K. R., de Vlugt, Y., Bramlage, A., Spaan, M., Elton, M., Snel, J., & Band, G. P. H. (2002). Alcohol consumption impairs detection of performance errors in mediofrontal cortex. Science (New York, N.Y)., 298(5601), 2209–2211. http://doi.org/10.1126/science.1076929

Romo, R., Hernández, A., & Zainos, A. (2004). Neuronal Correlates of a Perceptual Decision in Ventral Premotor Cortex. Neuron, 41(1), 165–173. http://doi.org/10.1016/S0896-6273(03)00817-1

Schultz, W., & Dickinson, A. (2000). Neuronal coding of prediction errors. Annual Review of Neuroscience, 23, 473–500. http://doi.org/10.1146/annurev.neuro.23.1.473

Ullsperger, M., & Danielmeier, C. (2016, February). Reducing Speed and Sight: How Adaptive Is Post-Error Slowing? Neuron. http://doi.org/10.1016/j.neuron.2016.01.035

Van der Borght, L., Desmet, C., & Notebaert, W. (2016). Strategy changes after errors improve performance. Frontiers in Psychology, 6(JAN), 2051. http://doi.org/10.3389/fpsyg.2015.02051

Van Der Borght, L., Schevernels, H., Burle, B., & Notebaert, W. (2016). Errors disrupt subsequent early attentional processes. PLoS ONE, 11(4),e0151843. http://doi.org/10.1371/journal.pone.0151843

Wallis, J. D., & Rushworth, M. F. S. (2013). Integrating Benefits and Costs in Decision Making. In Neuroeconomics: Decision Making and the Brain: Second Edition (pp. 411–433). http://doi.org/10.1016/B978-0-12-416008-8.00022-X

